# CoCoScore: Context-aware co-occurrence scoring for text mining applications using distant supervision

**DOI:** 10.1101/444398

**Authors:** Alexander Junge, Lars Juhl Jensen

## Abstract

Information extraction by mining the scientific literature is key to uncovering relations between biomedical entities. Most existing approaches based on natural language processing extract relations from single sentence-level co-mentions, ignoring co-occurrence statistics over the whole corpus. Existing approaches counting entity co-occurrences ignore the textual context of each co-occurrence. We propose a novel corpus-wide co-occurrence scoring approach to relation extraction that takes the textual context of each co-mention into account. Our method, called CoCoScore, scores the certainty of stating an association for each sentence that co-mentions two entities. CoCoScore is trained using distant supervision based on a gold-standard set of associations between entities of interest. Instead of requiring a manually annotated training corpus, co-mentions are labeled as positives/negatives according to their presence/absence in the gold standard. We show that CoCoScore outperforms previous approaches in identifying human disease–gene and tissue–gene associations as well as in identifying physical and functional protein–protein associations in different species. CoCoScore is a versatile text-mining tool to uncover pairwise associations via co-occurrence mining, within and beyond biomedical applications. CoCoScore is available at: https://github.com/JungeAlexander/cocoscore

## 1 Introduction

Text mining of the scholarly literature for the purpose of information extraction is a fruitful approach to keep abreast of recent research findings. The first step in information extraction is named entity recognition (NER) [Jurafsky and Martin, 2008]. Biomedical NER aims to identify relevant entities, such as genes, chemicals, or diseases, in text. Entities of interest can either be predefined in a dictionary or predicted using a machine learning model. NER is followed by a normalization step mapping the entities to a fixed set of identifiers, such as HGNC gene symbols [Yates et al., 2017] or Disease Ontology terms [Kibbe et al., 2015]. General approaches such as LINNAEUS [Gerner et al., 2010], Tagger [Pafilis et al., 2013], taggerOne [Leaman and Lu, 2016], or OGER [Basaldella et al., 2017] recognize diverse biomedical entities in text, while specialized tools recognize mentions of genetic variants [Allot et al., 2018] or chemicals [Jessop et al., 2011].

It is an active area of research to aggregate literature mentions of individual entities to extract higher-level information, such as pairwise biomedical relations, from the literature. Approaches to extract pairwise relations from a corpus of scientific articles, e.g., downloaded from PubMed, typically follow one of three main paradigms. Firstly, pattern-based approaches define a fixed set of regular expressions or linguistic patterns to match single phrases stating relations of interest. Pattern-based approaches typically achieve high precision but low recall in practice and require a labor-intensive manual construction of matching rules. Examples for this class of approaches are textpressso [Muller et al., 2004] or pattern-based approaches, as developed by Saric et al. [2004], used in STRING [Szklarczyk et al., 2017] and STITCH [Szklarczyk et al., 2016]. Secondly, unsupervised counting approaches count how often two entities appear together and aggregate these counts over the whole corpus in a co-occurrence statistic. A major shortcoming of simple counting-based co-occurrence scoring approaches to find pairwise relations is that the context of each co-occurrence is ignored, which can lead to low precision. For instance, sentences explicitly stating the absence of an association or describing findings unrelated to a relation are counted, too. Furthermore, counting-based co-occurrence scoring approaches do not allow to differentiate between different kinds of associations, such as physical protein–protein interactions and transcription factor–target associations. The major strengths of counting approaches are that they typically achieve relatively high recall and require no annotated training data or manually crafted match patterns. Examples of this class of approaches are the text-mining evidence contained in STRING and DISEASES [Pletscher-Frankild et al., 2015] as well as DisGeNet [Pinero et al., 2017]. Thirdly, supervised machine learning approaches require a labeled training dataset of associations and train a model to recognize relations of interest. Machine learning approaches are prone to overfit to the often small, manually annotated training datasets resulting in brittle models that do not generalize well to other datasets. For example Rios et al., 2018 showed that generalization between datasets of protein–protein and drug–drug interactions is only achieved when additional techniques such as the use of adversarial neural networks for domain adaption are employed. Examples for machine learning-based approaches to relation extraction are BeFree [Bravo et al., 2015], LocText [Cejuela et al., 2018], or conditional random fields [Bundschus et al., 2008].

Distant supervision [Mintz et al., 2009] is an approach similar to weak labeling [Morgan et al., 2004]. It can be used to generate datasets with a large number of samples with some amount of noise in the labels. Distant supervision for relation extraction only requires access to a knowledge base of well-described associations as well as an unlabeled set of entity co-occurrences. Labels for the dataset of co-occurrences are then inferred based on the presence or absence of the co-mentioned entities in the knowledge base. Note that a manually annotated text corpus is not required when using distant supervision.

In this work, we describe a novel approach, CoCoScore, that combines an unsupervised counting approach with a machine learning approach based on distant supervision. This allows CoCoScore to train a machine learning model to score sentence-level co-mentions without requiring an expert-curated dataset of phrases describing associations. The model is based on fastText [Joulin et al., 2016] and relies on word embeddings that represent words as dense vectors. CoCoScore finally aggregates all sentence-level scores in a given corpus in a final context-aware co-occurrence score for each entity pair. We apply CoCoScore to different biomedical relation extraction tasks: tissue–gene, disease–gene, physical protein–protein interactions, and functional protein–protein associations in *H. sapiens*, *D. melanogaster*, and *S. cerevisiae*. CoCoScore consistently outperforms a baseline model that uses constant sentence scores, following previously proposed approaches. We show then that the performance of CoCoScore further benefits from an unsupervised pretraining of the underlying word embeddings. By querying CoCoScore with manually constructed sentences, we show that keywords indicating protein–protein interactions and, to a certain extent, negations and modality are reflected in the sentence scores. A Python implementation of CoCoScore is available for download. The software package is geared towards reusability across many text mining tasks by only requiring a list of co-mentions for scoring without relying on a particular NER approach.

## 2 Materials and methods

### 2.1 Corpus

The corpus used for text mining consists of PubMed abstracts as well as both open access and author’s manuscript full text articles available from PMC in BioC XML format [Doğan et al., 2014, Comeau et al., 2018]. All abstracts were last updated on June 9th, 2018 and all full text articles were last updated on April 17th, 2018. We removed full text articles that were not classified as English-language articles by fast-Text [Joulin et al., 2016] using a pretrained language identification model for 176 languages downloaded from https://fasttext.cc/docs/en/language-identification.html. We furthermore removed full text articles that could not be mapped to a PubMed ID and those that mention more than 200 entities of any type included in our dictionary of biomedical entities such as proteins, chemicals, diseases, species or tissues. The final corpus consists of 28,546,040 articles of which 2,106,542 are available as full text and the remainder as abstracts.

### 2.2 Datasets and distant supervision

We use tagger v1.1 to recognize named entities in the corpus using a dictionary-based approach. Tagger can be downloaded from https://bitbucket.org/larsjuhljensen/tagger/. The dictionaries used for named entity recognition, training and test datasets as well as pretrained word embeddings and fastText scoring models described below can be downloaded from https://doi.org/10.6084/m9.figshare.7198280.v1.

The named entity recognition step is followed by a normalization step to a common naming scheme. All gene/protein identifiers were mapped to identifiers of corresponding proteins used in STRING v10.5 [Szklarczyk et al., 2017]. The normalization of disease and tissue identifiers is described below. We used placeholder tokens in all datasets to replace tissue, gene, protein, and disease names found by tagger. This blanking of entity names is important to learn a co-occurrence scoring model independent of the identity of the entities mentioned. Finally, we retain sentences that co-mention at least two biomedical entities of interest, depending on the given dataset.

The assignments of binary class labels to the sentences in each dataset follows a distant supervision approach to obtain a weak labeling. Given a sentence co-mentioning two entities of interest, the sentence is assigned a positive class label (1) if the entity pair is found in a given gold standard set of pairwise associations. If the two entities appear in the gold standard individually but not in association, the sentence is assigned a negative class label (0). The gold standard is specific to each dataset and described in the following sections. Table 1 lists information about the final datasets. Contrary to [Mintz et al., 2009], we treat each sentence in the dataset as a separate sample and do not merge all sentences for an entity pair into a single sample. This allows to define the final CoCoScore scoring scheme as a sum over all articles in the corpus (see Section 2.3).

**Table 1:** Entity, pair, and co-mention counts as well as as percentage of positive instances in all datasets.. ’Gold pairs’ refers to the total number of pairs found in the gold standard. ’Gold pairs cov.’ is the percentage of gold-standard pairs co-mentioned in at least one sentence in the dataset.

| Dataset type | Organism | Entity count | Pair count | Pos. pairs | Gold pairs | Gold pairs cov. | Co-mentions | Pos. co-mentions |
| --- | --- | --- | --- | --- | --- | --- | --- | --- |
| disease-gene | *H. sapiens* | 698 diseases, 1972 genes | 51,786 | 4.9% | 2,726 | 93.7% | 1,182,951 | 45.2% |
| tissue-gene | *H. sapiens* | 21 tissues, 14066 genes | 174,916 | 13.2% | 31,387 | 74.0% | 15,706,365 | 39.4% |
| functional PPA | *H. sapiens* | 4695 proteins | 1,032,063 | 14.1% | 361,744 | 40.4% | 16,390,304 | 58.2% |
| functional PPA | *D. melanogaster* | 1792 proteins | 36,524 | 27.4% | 88,604 | 11.3% | 358,141 | 64.9% |
| functional PPA | *S. cerevisiae* | 1567 proteins | 47,005 | 32.6% | 59,111 | 25.9% | 301,582 | 65.4% |
| binding PPI | *H. sapiens* | 6053 proteins | 1,236,751 | 2.9% | 76,299 | 47.4% | 15,611,378 | 21.3% |
| binding PPI | *D. melanogaster* | 2168 proteins | 60,378 | 12.7% | 32,541 | 23.6% | 463,822 | 42.6% |
| binding PPI | *S. cerevisiae* | 1612 proteins | 35,786 | 17.2% | 13,500 | 45.6% | 240,112 | 59.0% |

#### 2.2.1 Disease–gene associations

We followed the approach in the DISEASES database [Pletscher-Frankild et al., 2015] and obtained an expert-curated gold standard of disease–gene associations from GHR [Fomous et al., 2006] (downloaded on May 7th, 2017) by parsing disease-associated genes from json-formatted disease entries in GHR. We also retained entity co-mentions in the literature that involved a Disease Ontology (DO) [Kibbe et al., 2015] child term of a disease found in the gold standard. This propagation upwards the DO hierarchy yields a larger dataset of disease–gene associations while not compromising quality. Finally, disease names and aliases were mapped to DO identifiers.

#### 2.2.2 Tissue–gene associations

We followed the approach in the TISSUES database [Palasca et al., 2018] and downloaded manually curated tissue–gene associations from UniProtKB [SIB Members, 2016]. We restricted the tissue–gene association dataset to 21 major tissues, following the benchmarking scheme of the TISSUES database, and employed ontology propagation upwards the BRENDA Tissue Ontology (BTO) [Gremse et al., 2011], similar to the previously described DO propagation for disease mentions. Tissue names were normalized to BTO identifiers.

#### 2.2.3 Functional protein–protein associations

We obtained gold standard protein–protein associations (PPA) for *H. sapiens*, *D. melanogaster*, and *S. cerevisiae* following the approach for bench-marking associations in STRING [Szklarczyk et al., 2017]: Proteins found in at least one KEGG pathway map [Kanehisa et al., 2017] were considered positives since they are functionally associated in the given pathway. We also supplemented the original KEGG maps with artificial maps created by joining proteins from maps that share common metabolites.

#### 2.2.4 Physical protein–protein interactions

We obtained gold standard physical protein–protein interactions (PPI) for *H. sapiens*, *D. melanogaster*, and *S. cerevisiae* by obtaining interactions classified as ’binding’ from STRING v10.5 and retained only the highest scoring interactions, with a score > 0.9, as the gold standard. Binding interactions with a score ≤ 0.9, were added to a grey list. Co-mentions of grey-listed protein pairs were ignored and counted as neither positives nor negatives when creating the gold standard via distant supervision. While the resulting PPI datasets only contain protein pairs that physically bind to each other, the PPA datasets also encompass other functional associations as defined by membership in the same pathway.

### 2.3 Context-aware co-occurrence scoring

The context-aware co-occurrence scoring approach implemented in CoCoScore consists of two components. Firstly, a sentence-level classification model is trained to predict context-aware co-mention scores. Secondly, a scoring scheme combines sentence-level scores into a co-occurrence score taking evidence gathered over the whole corpus into account.

### 2.3.1 Unsupervised pretraining of word embeddings

Word embeddings represent each unique word in the corpus by a vector. We use a skipgram word embedding model that learns word vectors such that the vector representation of a word can be used to predict the words appearing in its context. This objective allows to represent words with similar syntax and semantics by similar vectors, as measured in terms of their inner product. Further details on the skipgram model and its training process can be found in Mikolov et al. [2013] and Bojanowski et al. [2016].

We pretrained word embeddings using fast-Text v1.0 [Joulin et al., 2016] on the whole corpus, not just on sentences in the dataset that co-mention entities of interest, which improves their generalization to downstream machine learning tasks. This step can be viewed as an instance of transfer learning where information is brought from a general task, the pretraining of word embeddings, to a specific task, the classification of sentences co-mentioning biomedical entities.

### 2.3.2 Training a sentence classification model

Our sentence-level classification model was implemented using fastText v.1.0 in supervised classification mode. Given a sentence, the pretrained vector representations of each word in the sentence are averaged. A logistic regression classifier *M* then predicts a binary class label since each sentence is labeled as either positive or negative after distant supervision. The sentence classification model *M* returns a score between 0 and 1. We interpret this score as a the probability that the sentence belongs to the positive class, i.e., that it states an associations.

We manually tuned the following hyperparameters of fastText: The dimensionality of word embeddings was set to 300; we performed 50 epochs of stochastic gradient descent with learning rate of 0.005 to train the model; we used unigram as well as bigram word features to partially capture local word order. Remaining hyperparameters were set to their defaults in the fastText v1.0 release.

### 2.3.3 Co-occurrence scoring

The final CoCoScore co-occurrence scores for a pair of entities aggregates the scores computed by sentence model *M* over all documents in the dataset.

Given a corpus *C* and an entity pair (*i, j*), the co-occurrence count *C*(*i, j*) for the pair is

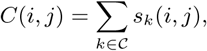

where

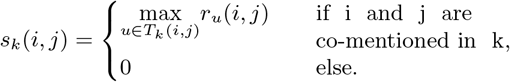

Here, *T_k_*(*i, j*) is the set of sentences co-mentioning *i* and *j* in document *k* and *r_u_*(*i, j*) is the sentence-level score returned by *M* for sentence *u*. The co-occurrence counts *C*(*i,j*) are converted to co-occurrence scores *S*(*i, j*) as follows:

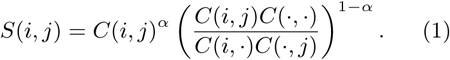

*C*(*i,*∙) and *C*(∙*, j*) are the sums of all co-occurrence counts involving entity *i* and *j*, respectively, *C*(∙,∙) sums the co-occurrences of all entity pairs. The hyperparameter *α* trades off the influence of *C*(*i, j*) counts and the observed-over-expected ratio captured in the second term of Equation 1.

Figure 1 outlines the complete context-aware co-occurrence scoring approach, illustrating both *C*(*i,j*) and *M*

**Figure 1:**
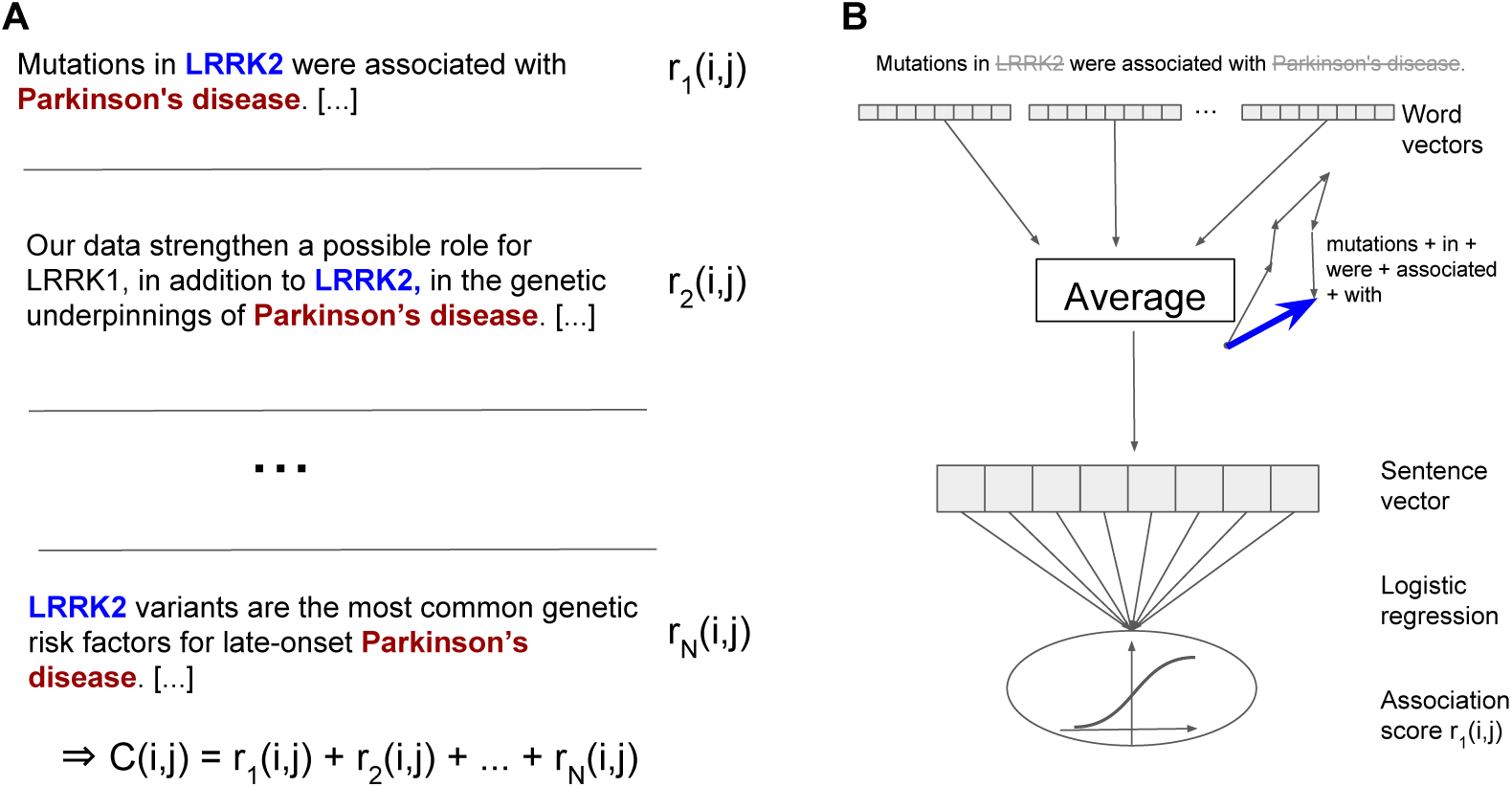
Cross-validation performance for CoCoScore and baseline. Values shown are the mean area under the precision-recall curve (AUPRC) across the three cross-validation sets. *α* chosen for each data set is depicted as vertical solid or broken lines, respectively. Note that AUPRC is not adjusted to a fixed prior since this adjustment does not affect the choice of *α*.

### 2.3.4 Baseline scoring scheme

We next defined a baseline model to compare CoCoScore to. Contrary to the context-aware model implemented in CoCoScore, the baseline model does not take context into account but scores all co-mentions equally. Given a corpus and entity pair (*i, j*), the baseline co-occurrence count *C̃*(*i, j*) is defined as:

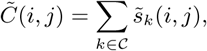

where

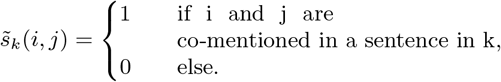

As before, the final co-occurrence scores *S*(*ĩ, j*) are computed from *C*(*ĩ, j*):

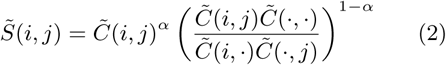

For the datasets of sentence-level co-mentions used in this work, this baseline model is equivalent to the co-occurrence scoring model used in, e.g. STRING [Franceschini et al., 2013, Szklarczyk et al., 2017], STITCH [Szklarczyk et al., 2016], TIS-SUES [Santos et al., 2015, Palasca et al., 2018], and DISEASES [Plestscher-Frankild et al., 2015].

### 2.3.5 Performance evaluation

We used the area under the precision–recall curve (AUPRC) to evaluate the co-occurrence scores. All AUPRC performance measures reported below were adjusted to a fixed percentage of 10% positive samples in the dataset. This adjustment makes AUPRC values comparable between datasets. We picked a positive percentage of 10% since this seems to be a realistic prior given our datasets Table 1.

The unadjusted AUPRC was computed by first sorting all pairs according to their co-occurrence scores in decreasing order and computing Precision and Recall at each score threshold as follows:

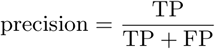

and

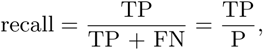

where TP is the number of true positives, FP is the number false of positives, FN is the number of false negatives, and P is the number of positives. The AUPRC is the area under the precision–recall curve. A random classifier has AUPRC equal to to the fraction of positives in the dataset and a perfect classifier has an AUPRC of 1. Precision–recall curves are better suited than receiver operating characteristic curves for dataset biased towards negatives since the latter give inflated performance estimates [Lichtnwalter and Chawla, 2012, Lever et al., 2016]. However, for comparison, we also state model performance in terms of area under ROC (AUROC) in the Supplementary Material of this article.

Adjusting the AUPRC to fixed class distribution was performed as follows: Let *a* be the aspired fraction of positives in the dataset (0.1 in this work) and *b* be the observed fraction of positives in the dataset. To adjust the AUPRC we replace Precision with the following adjusted measure:

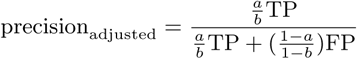

The adjusted AUPRC is then the area under the precision_adjusted_–recall curve.

### 2.3.6 CV and train/test splitting

For each dataset, we reserved all co-mentions involving 20% of the entity pairs as the test set which is only used for the final model evaluation step. This pair-level splitting ensures independence of training and test datasets since no entity pair is present in both training and test dataset. Each training dataset consists of co-mentions of the remaining 80% of entity pairs. In the training dataset, we randomly sampled a maximum of 100 sentence-level comentions per pair to ensure that the sentence-level model *M* does not overly fits to pairs that appear very often in the literature. To ensure a realistic performance evaluation, no such sampling was done for the test dataset. 3-fold cross-validation (CV) on the training set was used to tune the hyperparameter *α*. For computational reasons, we randomly sampled 10% of interactions in the three biggest datasets (functional PPA *H. sapiens*, binding PPI *H. sapiens*, tissue–gene associations) during CV. This reduced the number of associations in the downsampled dataset to approximately the number of associations in the remaining datasets.

## 3 Results and Discussion

### 3.1 Sentence scores of higher importance in CoCoScore than in baseline model

Before analyzing the performance on the test set, we tuned the weighting exponent hyperparameter *α* for both CoCoScore and the baseline model (see Section 2.3) via cross-validation (CV). *α* determines how much weight is put on the co-occurrence counts compared to the observed-over-expected ratio. The CoCoScore model achieved optimal CV performance for *α* ≈ 0.65 and the baseline model for *α* ≈ 0.55 for most datasets. Supplementary Figure 1 depicts CV performance depending on *α* for both models. We consider the tissue–gene dataset, where CV results in a considerably down-weighted observed-over-expected ratio term, an outlier due to the poor performance of both models on this dataset. The optimal *α* for CoCoScore was larger than the optimal *α* for the baseline model. This means that the best performing CoCoScore model put more weight on the co-occurrence counts than on the observed-over-expected-ratio, compared to the baseline model (Equations 1 and 2). We hypothesize that this is because CoCoScore down-weights uninformative sentences, compared to informative ones, making the sentence-level scores more reliable. Furthermore, the CoCoScore model outperformed the baseline on all datasets for the respective optimal *α* ranges. Below, we use *α* = 0.65 for CoCoScore and *α* = 0.55 for the baseline to compute test dataset performance.

### 3.2 CoCoScore outperforms baseline model in identifying disease–gene and tissue–gene associations

Table 2 lists AUPRC performance for both CoCoScore and the baseline model on the tissue-gene and disease-gene association datasets. Supplementary Table 1 depicts the performance in terms of the area under the receiver operating characteristic curve (AUROC). CoCoScore outperformed the baseline model on both dataset. Both approaches achieved considerably better performance on the disease–gene than on the tissue–gene association dataset. We thus manually inspected the 10 highest-scoring associations in the tissue–gene association dataset. Five of these tissue–gene pairs were counted as false positives, as defined by the gold standard derived from UniProtKB (Section 2.2.2). However, each of these pairs had more than 900 sentence-level co-mentions in articles and multiple sentences clearly stating the expression of the respective gene in the respective tissue. We concluded that these five association are likely true positives that are missing in the gold standard rather than rather than false positives. The seemingly poor performance on the tissue–gene association dataset can in part be explained by the incompleteness of the gold standard. While the low quality of the gold standard leads us to underestimate the performance, CoCoScore still appears to be able to learn informative text patterns leading to an improved performance.

**Table 2:** Adjusted area under the precision-recall curve (AUPRC) for CoCoScore and baseline model on tissue-gene and disease-gene association datasets generated via distant supervision.

| method | disease-gene | tissue-gene |
| --- | --- | --- |
| CoCoScore | 0.86 | 0.19 |
| baseline | 0.80 | 0.17 |

### 3.3 Physical protein–protein interactions are better identified than functional protein–protein associations

Figure 2 depicts performance on functional protein–protein associations (PPAs) and physical protein–protein interactions (PPIs) across *H. sapiens*, *D. melanogaster*, and *S. cerevisiae* for both CoCoScore and the baseline. Supplementary Figure 2 depicts the performance in terms of AUROC. CoCoScore outperformed the baseline and yielded similar improvements on all functional PPA and physical PPI datasets. While both models performed better on the binding PPI datasets than on the functional PPA datasets, we did not observe a clear trend in performance differences between organisms. CoCoScore achieves best adjusted AUPRC of 0.67 for binding PPI in *H. sapiens* and adjusted AUPRC of 0.57 in *D. melanogaster* and of 0.58 in *S. cerevisiae*. On the other hand, CoCoScore achieves best adjusted AUPRC of 0.50 for functional PPA in both *D. melanogaster* and *S. cerevisiae* and an adjusted AUPRC of 0.44 for *H. sapiens*. Overall, CoCoScore outperformed the baseline model on all six protein–protein association datasets surveyed.

**Figure 2:**
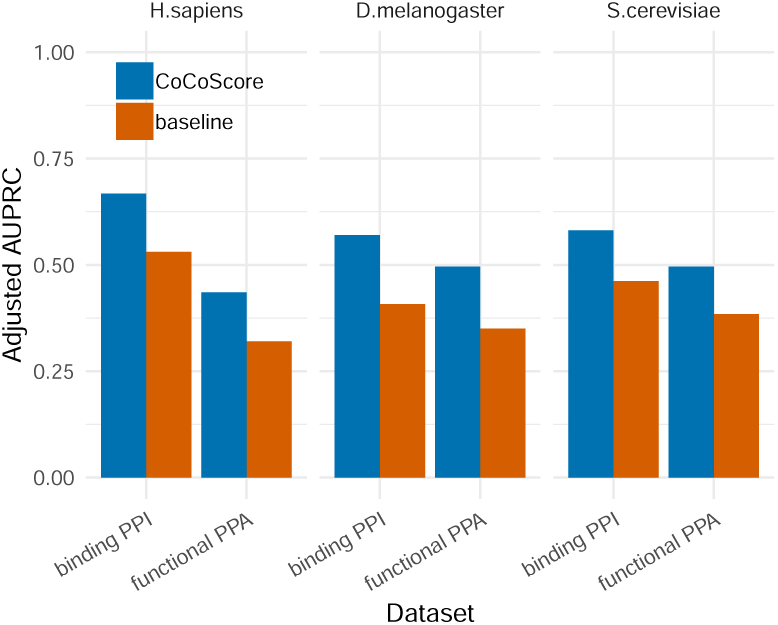
Performance for functional protein–protein associations and physical protein–protein interactions across *H. sapiens*, *D. melanogaster*, and *S. cerevisiae* for both CoCoScore (blue) and the baseline model (red). Performance is depicted as AUROC.

### 3.4 Pretrained word embeddings improve performance on most datasets

The default CoCoScore sentence classification model relies on word embeddings that were pretrained in an unsupervised manner on all articles in the corpus. To assess the impact of this pretraining step on CoCoScore’s performance, we compared the usage of pretrained word embeddings to the usage of embeddings that are learned along with weights for the logistic regression model at training time. Since the latter approach never accesses the complete corpus, word embeddings are only trained on sentences co-mentioning entities that are found in the respective training dataset.

Fig. 3 depicts adjusted AUPRC performance with and without pretrained word embeddings. The CoCoScore performance in Figure 3 is the same as shown in Figure 2 and Table 2. Supplementary Figure 3 depicts performance in terms of AUROC.

**Figure 3:**
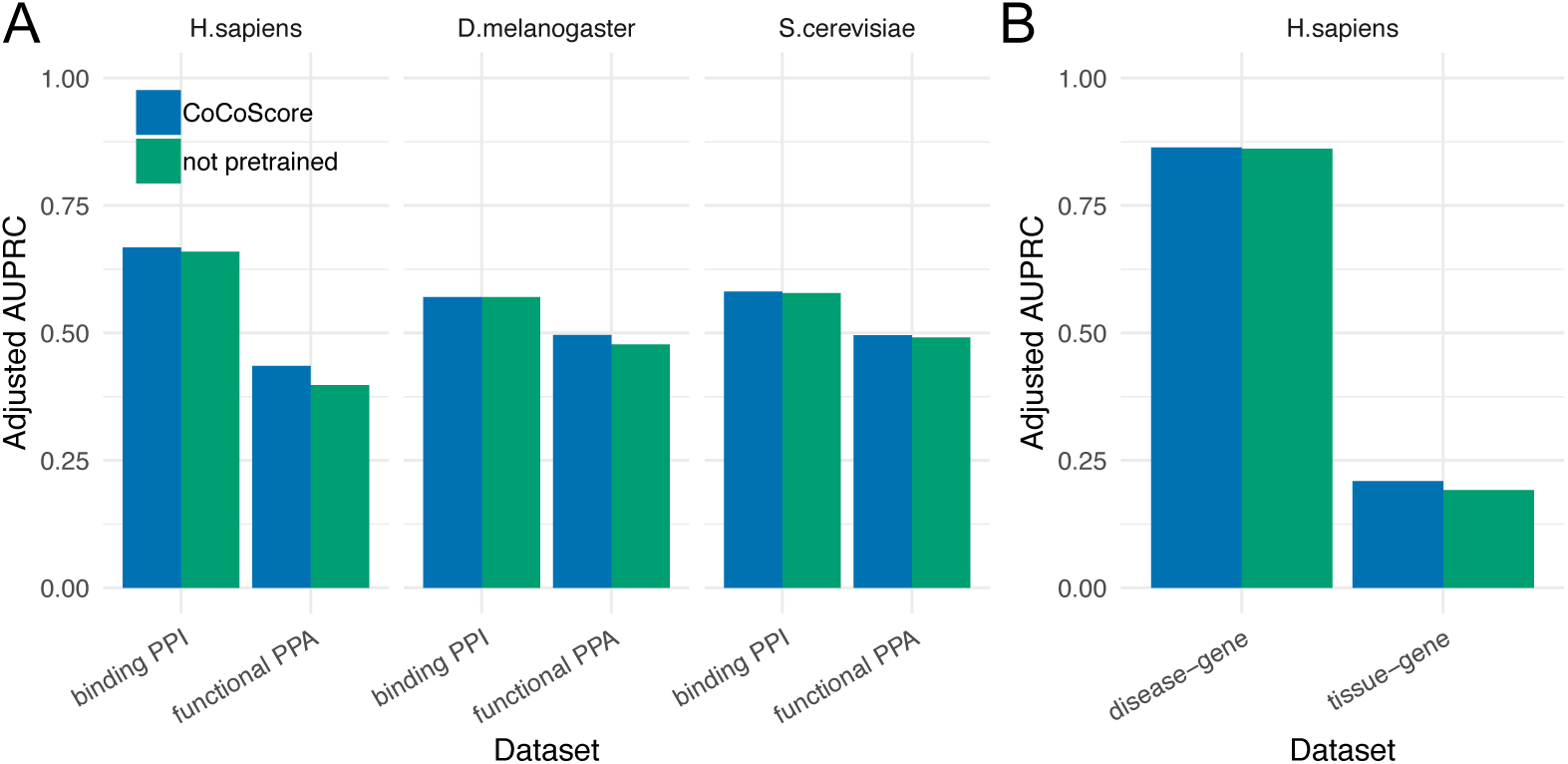
Performance with and without pretrained word embeddings for functional PPA and binding PPI datasts (A) as well as disease–gene and tissue–gene associations (B) for CoCoScore with (blue) and without (green) pre-trained word embeddings. Performance is depicted as AUROC.

The CoCoScore model using pretrained word embeddings in most cases outperformed non-pretrained word embeddings. We observe that the pretraining step was more fruitful for datasets with poor performance. The small increase in performance for some datasets could be due to the considerable size of the distantly supervised dataset (the smallest dataset contains 240k sentence co-mentions) which are large enough to train adequate word embeddings without pretraining. However, we still recommend using CoCoScore with pretrained embeddings for best performance.

### 3.5 Making sense of CoCoScore’s sentence scoring model by manually querying the model

The 300 dimensions of the word embeddings are not easily interpretable making it hard to understand which features drive sentence score predictions for a model trained on a given dataset. We thus used an indirect approach to interpret the sentence-level scoring model learned by CoCoScore by querying the model trained to recognize binding PPIs in *S. cerevisiae* with hand-crafted example sentences.

We observed that the model returned high scores for sentences containing keywords linked to physical interactions, such as the words ”complex” or ”sub-unit”, but did not pick up modality or uncertainty in sentences very well, once a keyword was present. For instance, the sentence ” and form a complex.” received a score of 0.99 while the sentences ” and do not form a complex.” and ” and might form a complex.” received a score of 0.98 and 0.99, respectively. Here, ” ” is a generic token used to blank protein names.

On the other hand, the model seemed to recognize negations and modality in sentences that contained the verb phrase ”bind to”. The sentence ” always binds to .” received a score of 0.72, ” binds to .” received a score of 0.44, ” possibly binds to .” received a score of 0.37, ” does not bind to .” received a score of 0.34, and ” never binds to .” received a score of 0.24. Based on this exploratory analysis, we conclude that the CoCoScore sentence scoring model for *S. cerevisiae* binding PPIs seems to rely on keywords and is able to detect modality and negations in certain situations.

### 3.6 Limitations and future work

While CoCoScore implements a novel context-aware co-occurrence scoring approach that improves upon a baseline model for all our test datasets, we see several limitations and directions for future research. Relations extracted by CoCoScore currently lack directionality. Many biomedical relations, such as protein phosphorylation, are directional that is not trivial to infer if, for instance, one protein kinase phosphorylates another, as commonly seen in signal transduction pathways. To address this shortcoming, CoCoScore’s distant supervision approach could be combined with pattern-based approaches to infer directionality. Alternatively, the word embeddings for the words in a sentence could not be collapsed into a single sentence vector but kept as a sequence of vectors fed into a sequence model such as a recurrent neural network.

We also plan to investigate the transferability of pretrained sentence scoring models between relation extraction tasks. For instance, a unified model could potentially be trained that recognizes not one specific type of relations, such as disease–gene associations, but also other relations, such as protein–protein interactions. Keywords and modality driving sentence scores (Section 3.5) should, to some extent, be transferrable between relation extraction tasks. Similarly, pretrained scoring models trained on one dataset could be combined with supervised learning performed on a second, expert-labeled dataset. This would enable the simultaneous use of large, distantly supervised dataset as well as small, accurately labeled dataset to boost performance. [Magge et al., 2018] use a similar approach to to identify geographic locations in sequence database entries.

Lastly, CoCoScore could be extended to score co-mentions beyond sentence-level by, for example, introducing a term in the scoring model that depends on the distance between entities co-mentioned outside a sentence.

## 4 Conclusion

Our newly developed approach, CoCoScore, performs pairwise co-occurrence scoring over a big corpus by combining an unsupervised counting scheme with a distantly supervised sentence scoring model based on pretrained word embeddings. The underlying sentence scoring model is able to recognize keywords, negations, and modality in sentences. Our approach performs better than a baseline scoring scheme inspired by previously proposed approaches on all eight benchmark datasets used in this study, covering disease–gene, tissue–gene, physical protein–protein interactions, and functional protein–protein associations.

CoCoScore is a versatile tool to aid biomedical relation extraction via text mining that is applicable to many applications beyond those presented in the paper. Our open source implementation only requires sentences co-mentioning entities as input and is available under a permissive license together with pretrained word embedding as well as the sentence scoring models trained in this work. This eases the integration of CoCoScore into existing text mining workflows for biomedical relation extraction.

## Acknowledgements

We would like to thank Rūdolfs Bērziņš for helpful discussions of this works as well as the team of the Danish National Supercomputer for Life Sciences (Computerome) for HPC support.

## Supplementary Material

**Figure 1:**
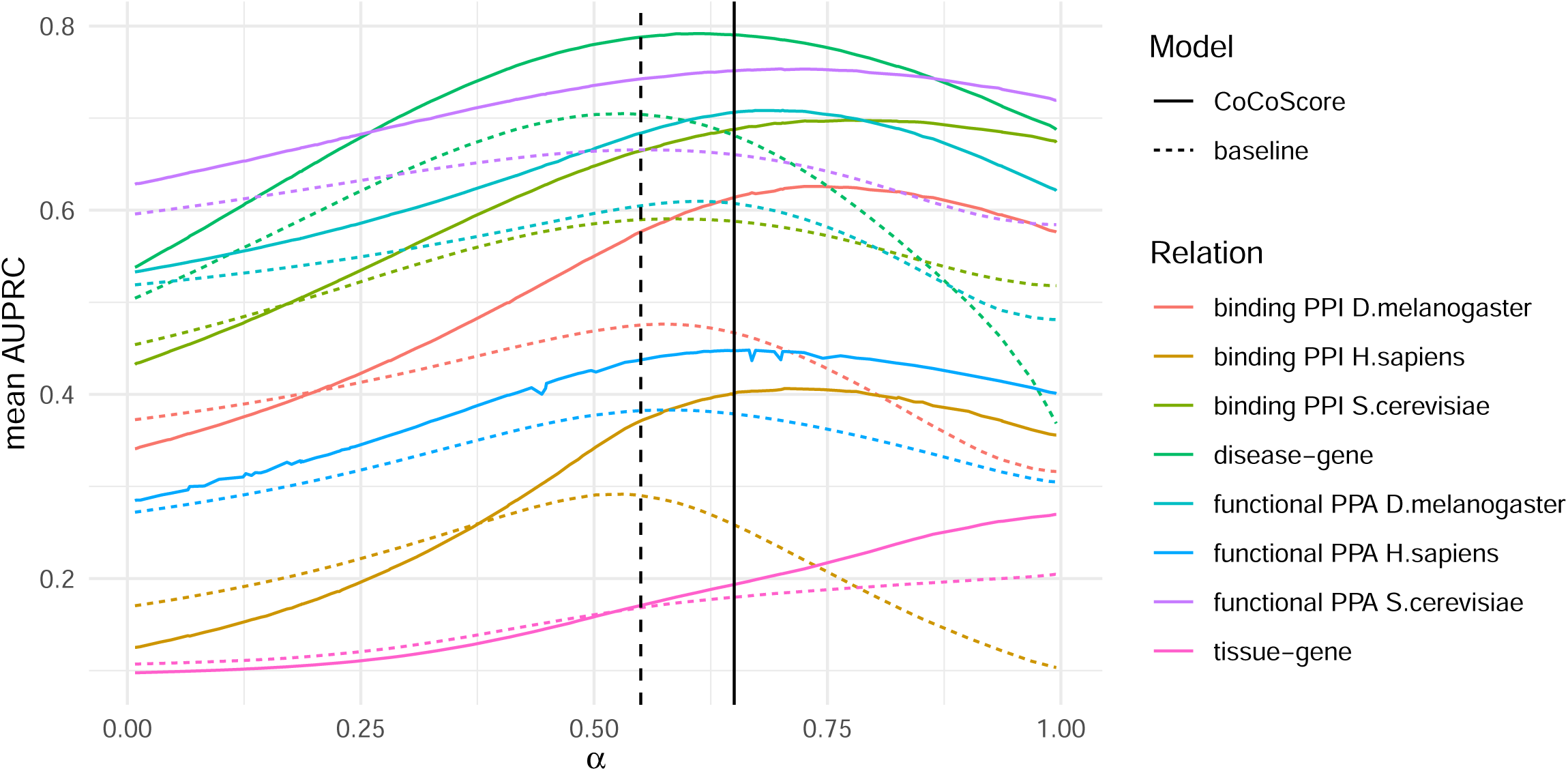
Context-aware scoring of co-occurrences. A) *N* sentences in the corpus co-mention the gene LRRK2 (*i*) and Parkinson’s disease (*j*). Context-aware sentence-level scores *r*(*i, j*) are summed to produce the final co-occurrence count *C*(*i, j*). B) The score *r*1(*i, j*) is computed by blanking gene and disease names, mapping all remaining words to their word vectors and scoring the resulting document vector with a logistic regression model (previously trained via distant supervision) (Section 2.2).

**Table 1:** AUROC for CoCoScore and baseline model on tissue-gene and disease-gene datasets generated via distant supervision.

| method | disease-gene | tissue-gene |
| --- | --- | --- |
| CoCoScore | 0.98 | 0.70 |
| baseline | 0.96 | 0.65 |

**Figure 2:**
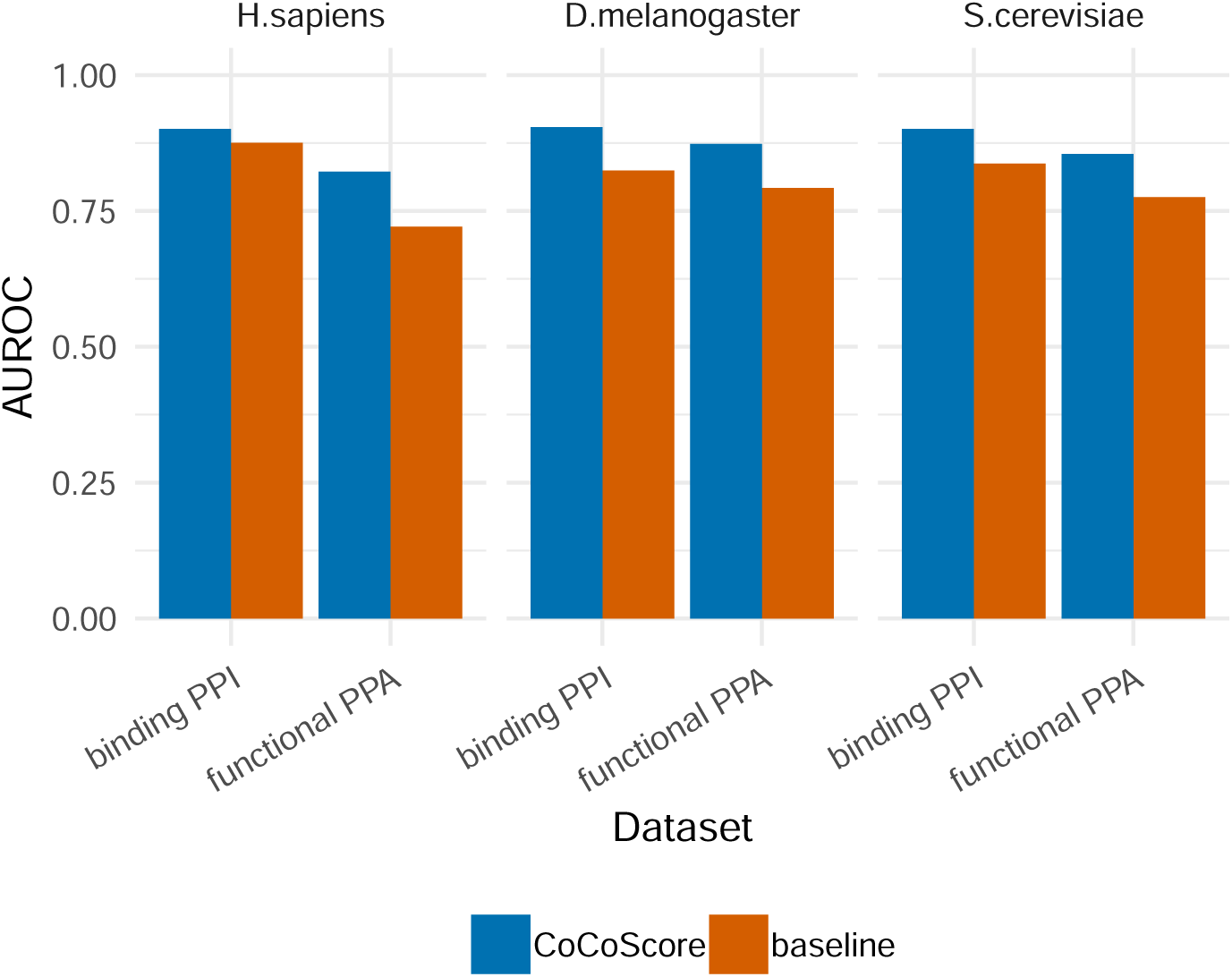
Performance on functional protein–protein associations and physical protein–protein interactions across *H. sapiens*, *D. melanogaster*, and *S. cerevisiae* for both CoCoScore (blue) and the baseline model (red). Performance is depicted as adjusted area under the precision-recall curve (AUPRC).

**Figure 3:**
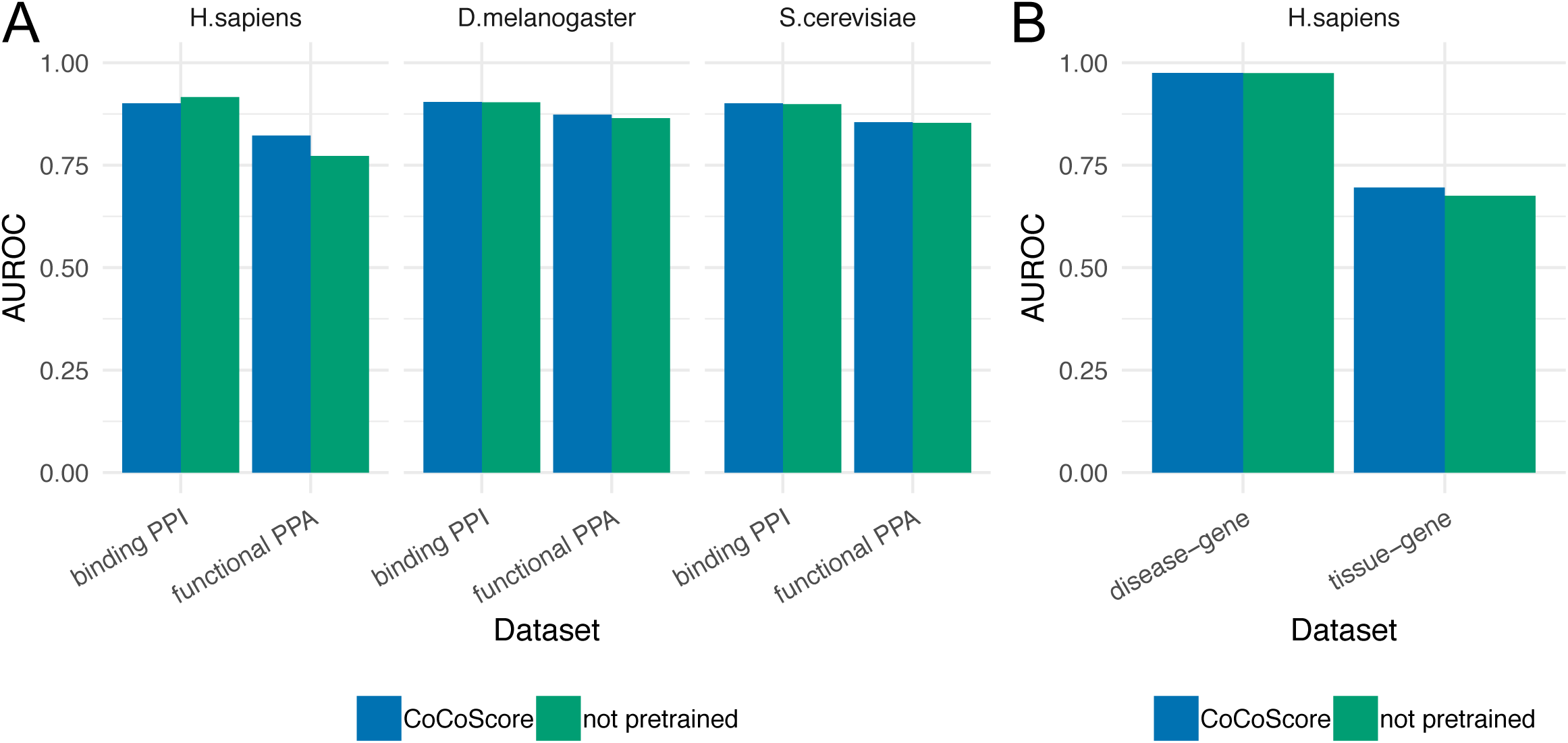
Performance with and without pretrained word embeddings on functional PPA and binding PPI datasets (A) as well as disease–gene and tissue–gene associations (B) for CoCoScore with (blue) and without (green) pretrained word embeddings. Performance is depicted as adjusted area under the precision-recall curve (AUPRC).).

